# Control of Male Mouse Copulatory Behavior by Midbrain Dopamine Neurons

**DOI:** 10.64898/2026.02.17.706366

**Authors:** Silvana C Araújo, Bertrand Lacoste, Luis Moreira, Oihane Horno, Eran Lottem, Joaquim Alves da Silva, Boris Gutkin, Christian Machens, Susana Q Lima

## Abstract

Ejaculation in male mice and humans requires intravaginal thrusting movements following penile intromission, yet the underlying neural mechanisms remain poorly understood. In this study, we used fiber photometry to record real-time activity of dopamine (DA) neurons in the ventral tegmental area (VTA) of male mice during copulation across six sexual interactions, in order to investigate how these dynamics change with experience and to examine their functional significance. Contrary to expectations, phasic VTA DA activity remained stable across sessions, showing time-locked responses to mounts, penile intromission, individual thrusts, and ejaculation, indicating experience-independent neural representations of all copulatory events. Notably, thrusting-related activity sharply declined just before ejaculation. Optogenetic inhibition of DA neurons disrupted thrusting behavior, but did not block ejaculation, suggesting DA signaling is critical for sustaining copulatory actions rather than triggering ejaculation itself. While prevailing theories frame VTA DA activity as primarily experience-dependent or centered on reward prediction, we here highlight its role in behavioral persistence during copulation. More broadly, our findings extend understanding of DA function beyond classical reward paradigms to the regulation of innate complex, goal-directed behaviors.

## Introduction

Sexual behavior is essential for species preservation and evolution. It consists of a coordinated sequence of actions that typically begins with pre-copulatory behaviors, such as approach and investigation of conspecifics (Lorenz 1981; Tinbergen 1951; Lenschow and Lima 2020). These behaviors increase sexual motivation and facilitate copulation in species with internal fertilization, providing genital stimulation that further enhances sexual arousal (Ågmo 2011; Lenschow and Lima 2020; Hull and Dominguez 2007; Brennan and Orbach 2020). Copulation culminates in the attainment of an “ejaculatory threshold,” at which point penile stimulation triggers ejaculation (Lucio et al. 2011).

In most mammals, copulation primarily relies on three male-driven actions: mounting (grasping the female’s flanks with the forelimbs), intromission (intravaginal penile insertion, often accompanied by thrusting), and ejaculation (sperm emission and expulsion) (Brennan and Orbach 2020; Lenschow and Lima 2020). While these elements are conserved, their temporal organization varies widely across species (Dewsbury 1972; 1975). For example, rats exhibit brief intromissions lasting 1–2 seconds followed by dismounting, whereas mice display prolonged intromissions lasting tens of seconds with continuous intravaginal thrusting (Dewsbury 1972; 1975; Lenschow et al. 2025; McGill 1962). The number of intromissions required to reach ejaculation also differs markedly, ranging from a single intromission in rabbits to multiple intromissions in rats and mice (Dewsbury 1972). Despite this interspecies diversity, copulatory behaviors are highly stereotyped within a species, reflecting evolutionary adaptations to specific ecological and social niches (Dewsbury 1972; Brennan and Orbach 2020; Eberhard 2019; Young and Wang 2004; Georgiadis et al. 2012; Pfaus et al. 2003). While the neural mechanisms underlying pre-copulatory behaviors have been extensively studied (Wei et al. 2021; Lenschow and Lima 2020), far less is known about how copulatory sequences themselves are encoded in the brain, how naïve animals can express them without prior experience, and how experience shapes sexual performance (but see Miyasaka et al. 2025; Bayless et al. 2023 for recent examples of studies).

Dopamine (DA) has long been implicated in human sexual behavior (Hyyppä et al. 2009), and early pharmacological studies demonstrated a central role for DA in multiple aspects of rodent sexual behavior (Malmnäs 1976; Tagliamonte et al. 1974; Pehek et al. 1988; Pfaus and Phillips 1989; Melis and Argiolas 1995; Mas et al. 1995). Subsequent microdialysis and voltammetry studies showed that dopaminergic input from the ventral tegmental area (VTA) to the nucleus accumbens (NAc) is released during both pre-copulatory and copulatory phases of sexual behavior (Pleim et al. 1990; Pfaus et al. 1990; Wenkstern et al. 1993; Damsma et al. 1992; Fumero et al. 1994; Robinson et al. 2002; 2001; Beny-Shefer et al. 2017). More recently, genetically encoded tools have revealed that VTA dopaminergic neurons are closely associated with approaching social stimuli (Gunaydin et al. 2014; Solié et al. 2022) and that DA is released in the NAc during mounting, thrusting, and ejaculation in male mice (Sun et al. 2020; Dai et al. 2022; Miyasaka et al. 2025).

The initial debate whether VTA DA neurons encode reward prediction errors (important for error-driven learning (Schultz et al. 1997)), or represent the incentive salience of natural and learned stimuli (fundamental for motivation and the immediate control of behavior (Robinson 1993)), has progressively expanded. VTA DA has also been involved in the effort dedicated to obtaining motivationally relevant stimuli (Salamone et al. 2003; Hamid et al. 2016), performance (Bakhurin et al. 2023), learning rate (Coddington et al. 2023), reward itself (Wise and Rompre 1989), and in the encoding of sensory, cognitive and movement variables (Engelhard et al. 2019). Notably, all of these processes are engaged during copulation, complicating functional interpretations.

Here, we used fiber photometry and optogenetic manipulations to examine the activity of VTA DA neurons in male mice across their first six copulatory experiences. Despite variability across animals and sessions in the number of intromissions required to reach ejaculation, population-level analyses revealed no systematic changes in the total number or duration of intromissions with repeated experience, indicating that repeated cycles of mounting, intromission, and intravaginal thrusting represent a core feature of the mouse reproductive strategy. From the very first copulatory encounter, VTA DA neurons exhibited transient activation during mounting, individual thrusts, and ejaculation, a pattern that remained stable across experience. Strikingly, arrival at the ejaculatory threshold was marked by an increase in thrusting rate during the final mount and a concomitant decrease in VTA DA activity, suggesting that DA signals reflect the animal’s internal state. Transient inhibition of VTA DA neurons during approach suppressed mounting, and inhibition after mount initiation disrupted thrusting, without preventing ejaculation itself. Together, these findings indicate that VTA DA activity provides a feedback signal necessary for sustaining mounting and thrusting behaviors until the ejaculatory threshold is reached.

## Results

### Neural dynamics during sexual interaction

We allowed ten sexually naïve male mice to mate with sexually experienced, ovariectomized, and hormonally primed females (Lenschow et al. 2025; Valente et al. 2021) until each male reached 6 ejaculations (Fig. 1A, 6 sessions, see Methods). To monitor population activity of ventral tegmental area (VTA) dopamine (DA) neurons during copulation, we performed fiber photometry recordings (Matias et al. 2017), to measure calcium-dependent activity (Fig. 1B). Briefly, we unilaterally injected an adeno-associated virus (AAV) encoding a Cre-dependent calcium indicator (GCaMP6f) or yellow fluorescent protein (YFP) control (Addgene), driven by the synapsin promoter, into the VTA of transgenic mice expressing Cre recombinase under the dopamine transporter (DAT) promoter (Bäckman et al. 2006). The DAT-Cre line has previously been shown to selectively label VTA dopamine neurons (Lammel et al. 2015). Male mice were additionally implanted with a single optical fiber positioned above the VTA (Matias et al. 2017). Fiber placement and viral expression were verified histologically at the end of the experiment (Fig. 1C, see Methods).

**Figure 1.**
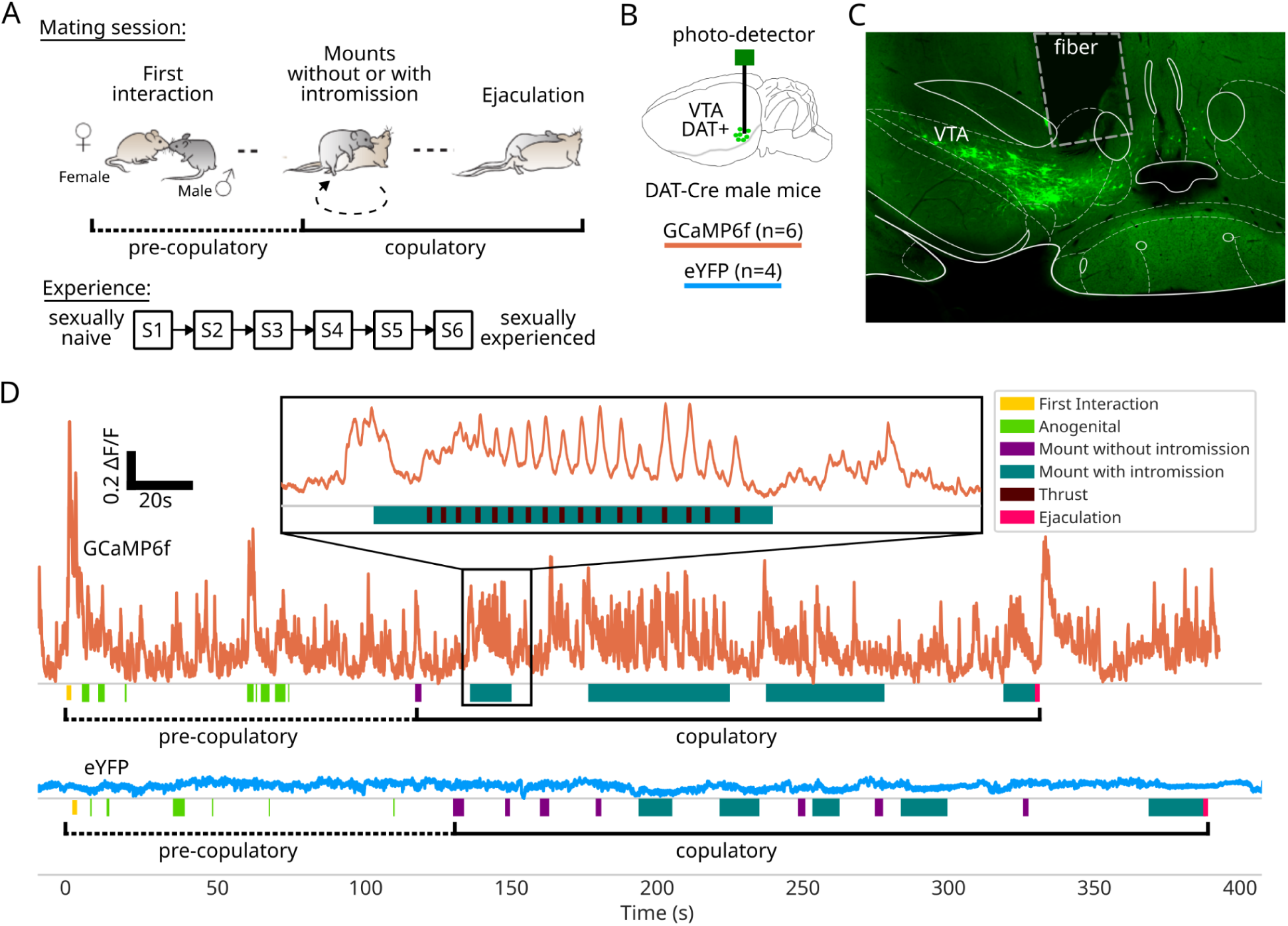
Neural dynamics during sexual interaction. (A) Schematic of mice sexual behavior, with a pre-copulatory phase containing mutual investigations, especially anogenital investigations, and a copulatory phase containing Mounts with and without penile insertion, intra-vaginal thrusts, and ejaculation. (B) Fiber implantation in the VTA of DAT Cre males with a GCaMP6f (n=6) or YFP (n=4). (C) Representative image showing GCaMP6f expression and fiber placement in the VTA. (D) Two example sessions from two animals expressing GCaMP (Top, session 3) and YFP (Bottom, session 5). For each session, the (not-normalized) ΔF/F is shown above a raster of the annotated behavioral events We distinguish the two phases: pre-copulatory and copulatory. Inset: zoom on the first mount of the GCaMP session, showing the ΔF/F with the peaks associated with the Thrusts shown on the raster below. X-axis is zoomed by x10, while Y-axis scale is identical.

Following an initial pre-copulatory phase characterized by investigatory behaviors, most prominently anogenital investigations, male mice attempted copulation by grasping the female with their forepaws (Fig. 1A). These attempts either resulted in penile insertion (mount with intromission, MI) or did not (mount without intromission, MwoI). MwoIs were typically brief (1-2 s), whereas MIs were longer-lasting events (10-20 s) accompanied by rhythmic pelvic thrusting with continuous vaginal penetration (Lenschow et al. 2025; McGill 1962; Dewsbury 1975). If ejaculation did not occur, males dismounted the female and repeated a variable number of mount–intromission-dismount sequences until ejaculation was achieved. Ejaculation was readily identifiable by a characteristic shivering response followed by the male falling to the side of the female and remaining immobile for several seconds before separation (Lenschow et al. 2025; Valente et al. 2021). Although some sessions were aborted before mounting occurred, in all sessions in which copulation was initiated males ultimately reached ejaculation (Supplementary Table 1; see Methods). We manually annotated the first interaction event, all anogenital investigations, and all copulatory behaviors (MI, MwoI, thrusts, and ejaculation) across all six sessions for each male.

Figure 1D shows representative photometry traces from two sessions recorded in two different males: a GCaMP6f-expressing male (top; inset shows a single mount) and a YFP control male (bottom). We observed pronounced increases in VTA DA neuron activity during sexual interactions in GCaMP6f mice, whereas no such transients were present in YFP controls, indicating that these signals reflect calcium-dependent neural activity rather than motion artifacts. With repeated sexual experience, we observed modest reductions in the duration of the pre-copulatory phase (time from first interaction to first MI or MwoI, Supp. Fig. 1A) and overall copulatory duration (time from first MI or MwoI to ejaculation, mainly due to an overall modest decrease in the intermount intervals across time, Supp. Fig. 1B), but no further changes in behavioral performance. Specifically, the number of MIs, MI duration, intermount interval (time interval between two consecutive MIs) and fraction of MIs remained stable across sessions (Supp. Fig. 1C-F). Together, these results indicate that copulatory behavior remained largely invariant across repeated sexual experiences.

### VTA DA activity during the first interaction, anogenital investigations and ejaculation as a function of experience

We started by investigating the nature of the recorded transients at the boundaries of the sexual interaction: the first interaction and anogenital investigations, and ejaculation. The first interaction (first time the animals interact after the female enters the behavioral box) with the female was the behavioral event accompanied by the largest change in fluorescence in every session (Supp. Fig. 2A). Since the activities (dF/F) are different across animals, we normalized the activity of VTA DA neurons in order to average over animals. Specifically, we used the peak fluorescence of the first interaction to normalize fluorescence changes across the rest of the session (see Methods). Using the average signal change accompanying MwoI/MI as a normalization factor for each session leads to similar values to the ones obtained when using the amplitude of the first interaction (Supp. Fig. 2A). Normalized fluorescence signals could then be pooled across different sessions and animals. The temporal dynamics of the fluorescence signal observed during the first interaction was similar between the first and last sessions (Fig. 2A). In contrast, while anogenital investigations during the first session were accompanied by a small change in fluorescence, with experience those transients disappeared (Fig. 2B). Ejaculation was also accompanied by a large transient that persisted for several seconds (Fig. 2C). Surprisingly, even though the males became progressively experienced, we observed no difference in the activity accompanying ejaculation when comparing session 1 with session 6 (Fig. 2C) or when comparing the peak activity across all the sessions (Fig. 2D). The peak activity observed during ejaculation was not related to the pre-copulatory phase (Fig. 2E), the copulatory phase duration (Fig. 2F), the number of MIs (Fig. 2G), nor the total number of Thrusts (Fig. 2H).

**Figure 2.**
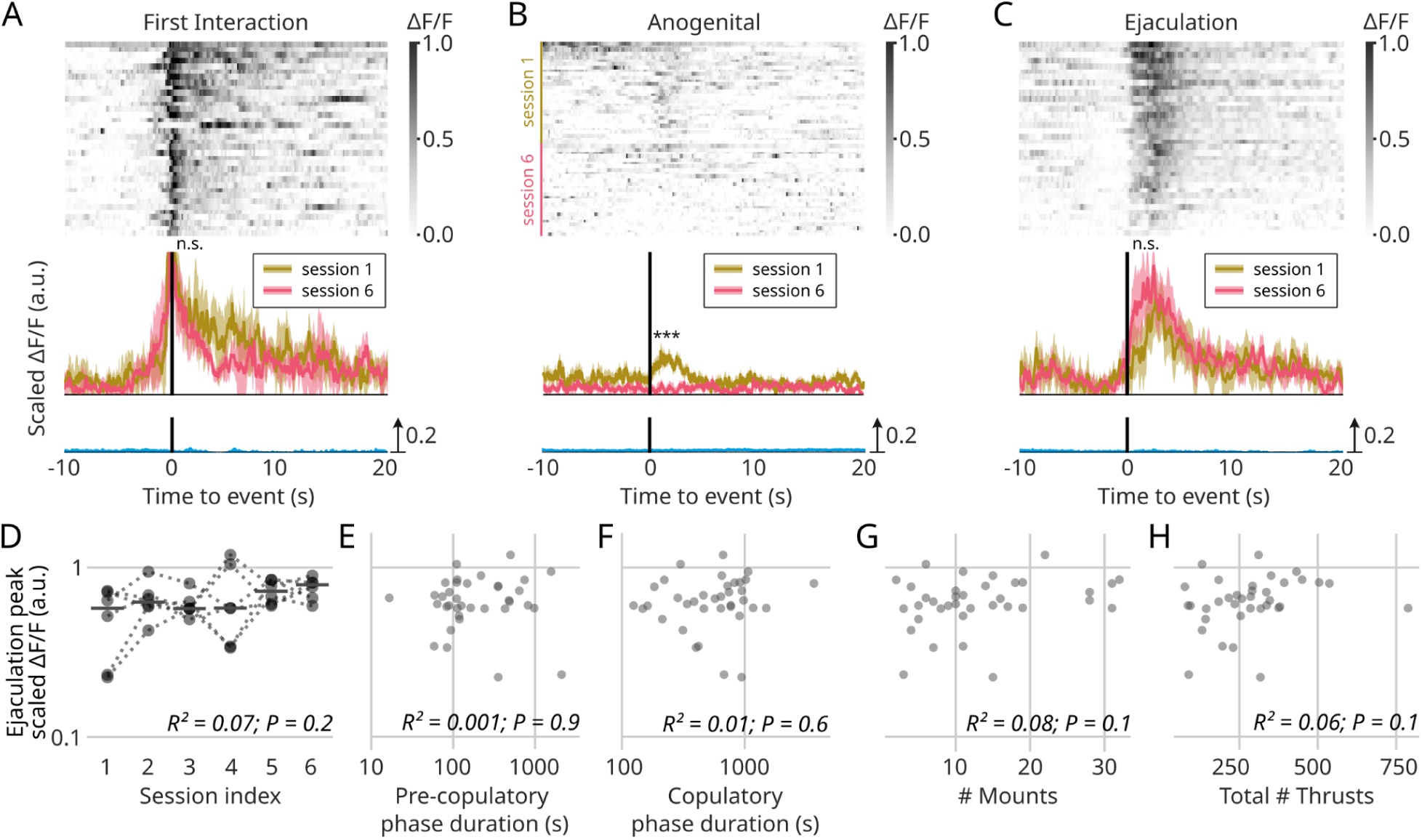
VTA DA activity during the first interaction, anogenital investigations and ejaculation as a function of experience. (A-C) (Top) Raster of the scaled ΔF/F for the GCaMP sessions, aligned to different events: (A) First interaction, (B) Anogenital investigation and (C) Ejaculation. For First interaction and Ejaculation, the rasters contain all the GCaMP sessions, whereas for Anogenital, only the first and the last sessions. (Bottom) Median scaled ΔF/F aligned to the event for YFP sessions (bottom, blue); for the first session, naive session, of GCaMP animals (top, yellow); for the last experienced session of GCaMP animals (top, pink). Standard error of the median as shaded error ribbons. Comparison of the average signal up to 2s after each event for GCaMP sessions between the first and the last sessions, two-tailed Student’s t-test: First interaction, p = 0.4; Anogenital, p = 0.0009; Ejaculation, p = 0.1. (D-H) Ejaculation peak (scaled ΔF/F for the GCaMP sessions) in log-scale for all animals: (D) across sessions, (E) in relationship to the pre-copulatory phase duration (in log-scale), (F) to the copulatory phase duration (in log-scale), (G) to the number of mounts before ejaculation and (H) to the total number of Thrusts performed to reach ejaculation. (D) Linear regression R² and t-test for the slope. (E-F) Pearson’s R² and test for the null hypothesis that the correlation is zero, for each type of mount.

In contrast to all other annotated behaviors, which were not accompanied by fluorescence changes in the YFP animals, genital grooming was accompanied by a modest change in fluorescence that was detected both in animals expressing the calcium indicator and YFP alone, suggesting that fluorescence changes in this case were due to movement artifacts (Supp. Fig. 2E).

### The activity of VTA DA neurons associated with mounting does not change with sexual experience or session progression

We observed an increase in activity that started ramping up before the male placed his two front limbs on the female’s back, and which was similar between MwoI and MI (Fig. 3A,B). Given that our mice were allowed to copulate six times, progressing from sexually naive to sexually experienced, we wondered if the mount-associated signal evolved with experience. In line with the lack of changes in the behavioral performance of the males, the population activity of VTA DA neurons accompanying each mount event was similar between sexually naive (Session 1) and sexually experienced males (Session 6) (Fig. 3A-C).

**Figure 3.**
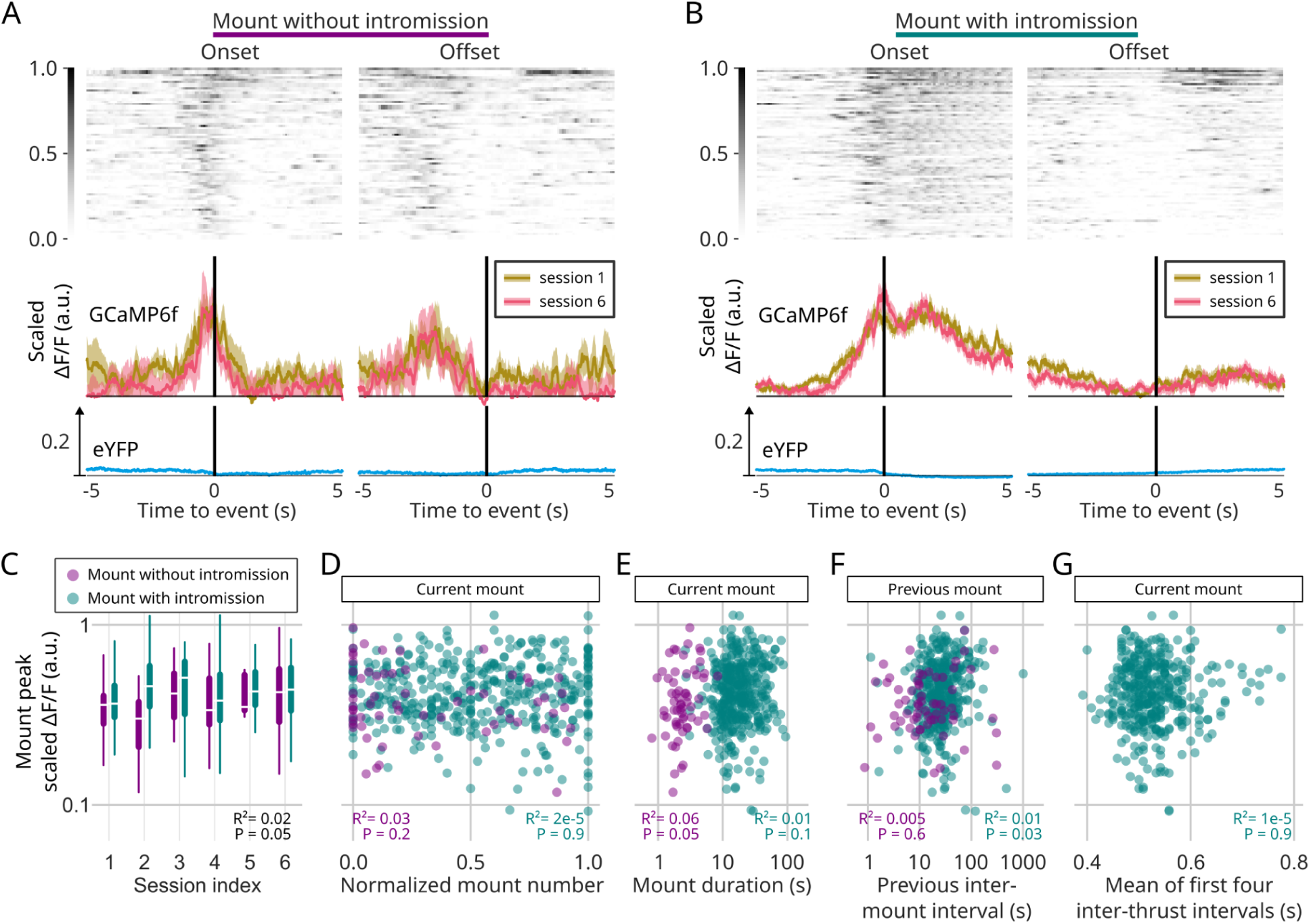
The activity of VTA DA neurons associated with Mounts does not change with sexual experience or session progression. (A-B) (Top) Raster of the scaled ΔF/F for the GCaMP sessions, aligned to different events: Mount onset and offset, split between Mounts without intromission (A) and with intromission (B). The scaled ΔF/F amplitude is from 0 to 1. (Bottom) Median scaled ΔF/F aligned to the event for YFP sessions (bottom, blue); for the first session, naive session, of GCaMP animals (top, yellow); the last experienced session of GCaMP animals (top, pink). Standard error of the median as shaded error ribbons. (C-G) Mount peak (scaled ΔF/F for the GCaMP sessions) in log-scale for all animals across: (C) sessions; (D) normalized mount number; (E) Mount duration (in log-scale); (F) previous inter-mount interval (in log-scale). (G) mean of the first four inter-thrust intervals (only for Mounts with Intromission). Dot color refers to mount identity, Mount without intromission (purple) or Mount with intromission. For C, D, E and G, it refers to the identity of the current mount, whose peak is measured. For F, it refers to the identity of the previous Mount. (C) Linear regression R² and t-test for the slope for both types of mounts together. (D-G) Pearson’s R² and test for the null hypothesis that the correlation is zero.

The activity associated with MwoI and MI was also not related to the position of the mount within the session (Fig. 3D), the current mount duration (Fig. 3E), the previous mount interval (the interval between consecutive mounts, Fig. 3F), nor the vigor of the first 4 thrusts (average interval between the first four thrusts after MI (Fig. 3G).

### The activity of VTA DA neurons associated with thrusts does not change with sexual experience, session progression or vigor

Finally, we focused on the VTA-DA GcaMP signal around the thrust events. Sometimes during a MI, the male might stop thrusting without dismounting the female, reinitiating thrusting after a couple of seconds with the initial vigor (backpacking events, see Methods and Supp Fig. 3A for an example). Because of that, we considered only the thrusts belonging to the first thrusting bout, which ends when an inter-thrust interval lasts more than 1.2s. Each individual thrust was accompanied by a short burst in activity (Fig. 4A,B), whose amplitude did not change with experience (Fig. 4C) and was not present in the YFP controls (Fig. 4B). Finally, even though animals thrust at a stereotypical frequency, there is variability on the exact timing of each thrust. We found a significant, although not sizable, correlation between the thrust peak amplitudes and the previous inter-thrust interval (Fig. 4D). We also observed a significant, but not sizable, correlation between the thrust peak and the following inter-thrust interval (Supp Fig. 3E). We can also quantify the difference or error between the real and expected inter-thrust interval (see Methods). We found a barely significant, not sizable, correlation between the error and thrust peak amplitude (Supp Fig. 3F).

**Figure 4.**
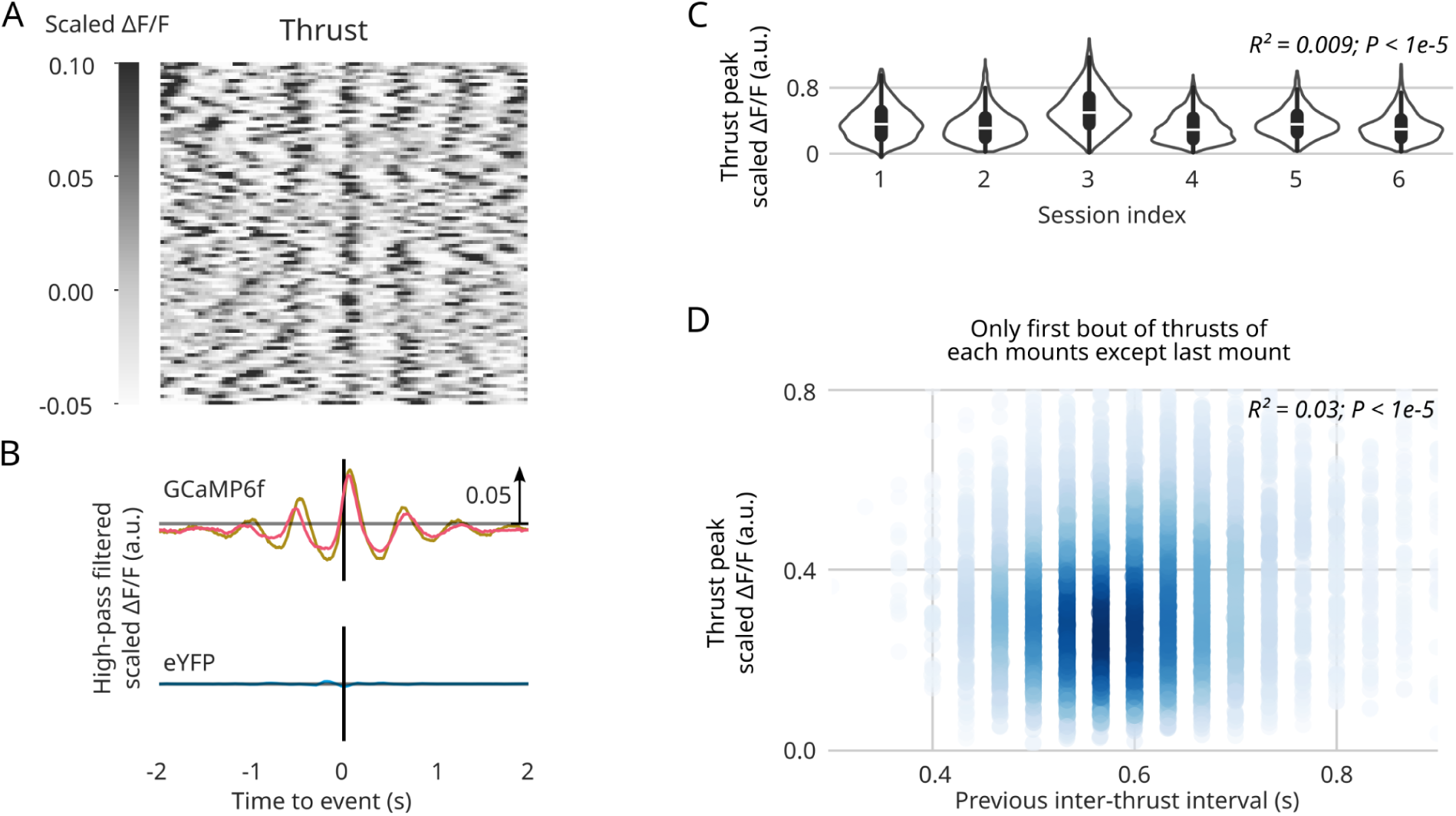
The activity of VTA DA neurons associated with Thrusts does not change with sexual experience, session progression or vigor. (A) Raster of the scaled ΔF/F (high-pass filtered) for the GCaMP sessions, aligned to individual Thrusts. The scaled ΔF/F amplitude from -0.05 to 0.1. (B) Median scaled ΔF/F (high-pass filtered) aligned to the event for YFP sessions (bottom, blue); for the first session, naive session, of GCaMP animals (top, yellow); the last experienced session of GCaMP animals (top, pink). Standard error of the median as shaded error ribbons. (C) Thrust peak (scaled ΔF/F, high-pass filtered) versus session index for all animals (except last mount). Linear regression R² and t-test for the slope. (D) Thrust peak (scaled ΔF/F, high-pass filtered) versus previous inter-Thrust interval. Pearson’s R² and test for the null hypothesis that the correlation is zero.

In summary, all copulatory events (MwoI, MI, thrusts and ejaculation) are accompanied by increases in fluorescence with different amplitudes and temporal dynamics that do not change across a session nor with experience.

VTA DA neurons are known to be active when the animal is receiving a reward or, after a learning phase, when a rewarding-predicting time-locked cue is presented before the reward (Schultz et al. 1997). Even though we observed a decrease in the anogenital locked activity with experience (Fig. 2B), we were surprised with the lack of experience dependent changes in the activity of VTA DA neurons during the rest of the annotated behaviors. Because of that, we reproduced a Pavlovian conditioning experiment with two other males expressing GCaMP6f in VTA DA neurons (Supp. Fig. 4A). Fluorescence changes were recorded during a simple cue-reward task, without normalizing the ΔF/F by animal. When the animals started the task, we immediately observed a peak of the GCaMPf signal after receiving the liquid reward associated with licking (Supp Fig. 4B, top). After the animal learned the association between a visual cue and the subsequent liquid reward, we observed a peak of activity time-locked to the cue start (Supp Fig. 4B, bottom). Then we recorded the activity of VTA DA neurons while these animals copulated (2-4 copulation sessions). We found that the amplitude of the signal around the First interaction and Ejaculation was similar to the amplitude of the signal at the cue and liquid reward, with smaller Mounting and Thrusts signals (Supp Fig4 C). So, the recorded VTA DA neurons behave as expected in the context of a Pavlovian task, this is, the activity retrogradely moves to the neutral cue that predicts the reward. In contrast, the relationship between the activity observed at the first interaction and the activity observed during copulatory events (mounts, intromission and ejaculation) did not change with experience, suggesting these are not artifacts, but rather real signals.

### Behavior and neuronal activity changes in the last mount

In the previous sections, the last mount before ejaculation (Fig. 5A) was excluded from the analysis because it clearly differed from all the other mounts. During all MIs (except the last) the thrusting rate decreases until the male dismounts the female (Fig. 5B, see also McGill 1962). The last MI before ejaculation, however, starts with a slightly higher thrusting rate which then accelerates (Fig. 5B). This acceleration of the thrusting rate close to ejaculation is present from the first session (Fig. 5B). Despite these behavioral changes, the peak DA activity during the mount onset is similar between the last mount and all other mounts (Fig. 5C).

**Figure 5.**
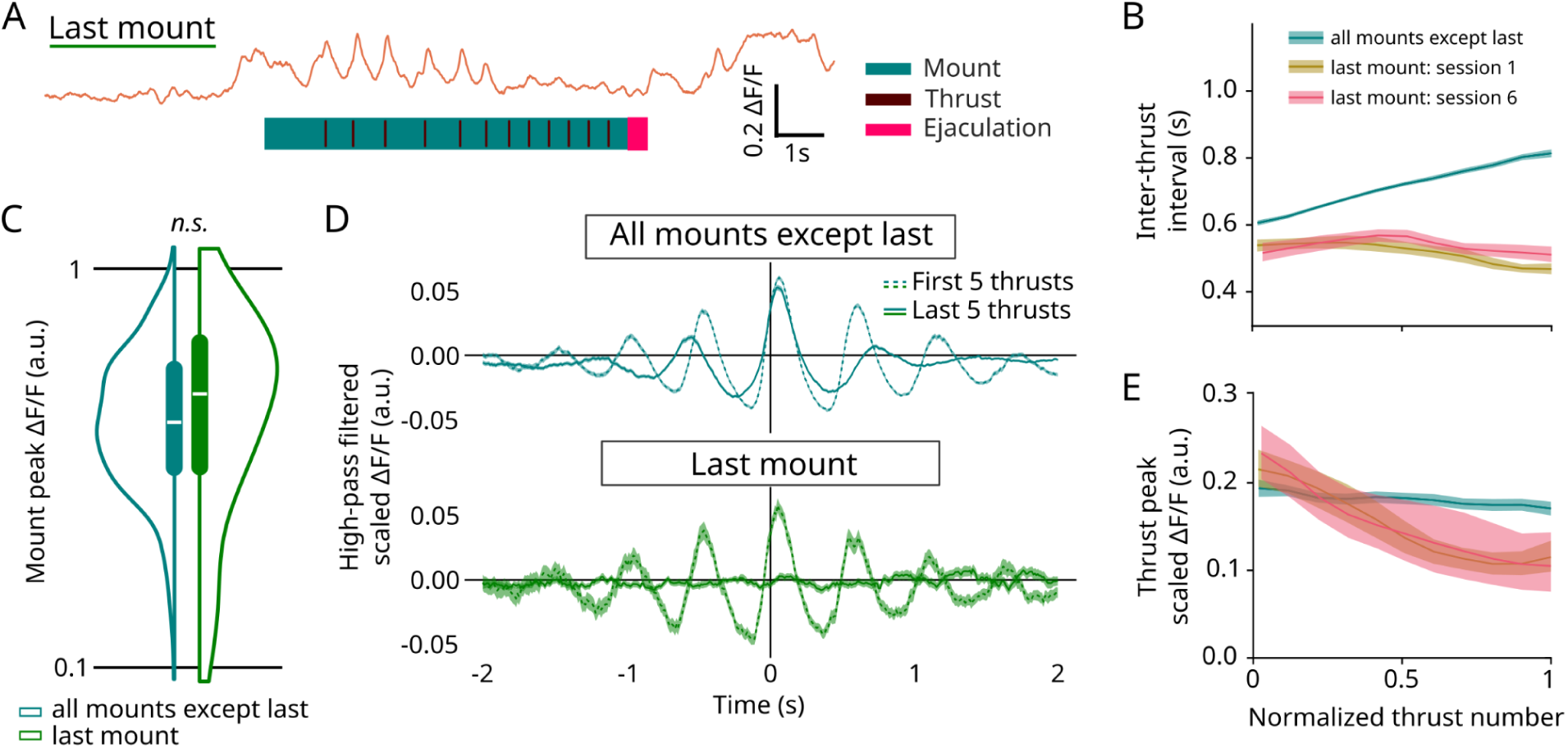
Behavior and GCaMP changes in the last mount. (A) One example of the (not-scaled) ΔF/F in the last mount (first session) above a raster of the annotated behavioral events. (B) Inter-thrust interval versus normalized thrust number only for the first thrusting bout (that is the thrust index in the mount normalized by the number of thrusts in the first bout) for thrusts in the last mount of the first session, naive session (yellow); in the last mount of the last experienced session (pink); in all the mounts (all sessions) except the last mount (cyan). Conditional standard error of the mean as shaded error ribbons. (C) Distribution of Mount peaks (scaled ΔF/F for the GCaMP sessions, Mounts with Intromission) in log-scale for all animals across sessions, split between all mounts except last (cyan) and the last mounts (green). Two-tailed Student’s t-test p = 0.1. (D) Median scaled ΔF/F (high-pass filtered) aligned to Thrust events, for the first 5 thrusts of the Mounts (dotted line) and the last 5 thrusts of the Mounts (solid line): (Top) for all the mounts except the last mount of each session; (Bottom) for the last mount of each session. (E) Same as (B) but for the Thrust peak (scaled ΔF/F, high-pass filtered) versus normalized thrust number only for the first thrusting bout.

Interestingly, the increase in thrusting rhythm of the last mount is associated with a disappearance of the activity in VTA DA neurons (Fig. 5D, E). This is illustrated by the PETH aligned to the five first thrusts of a MI versus the last five thrusts of a MI, for the other mounts (Fig. 5D top) and for the last MI before ejaculation (Fig. 5D bottom). These PETHs exhibit a peak around the thrust event, in all cases except for the last thrusts of the last MI, revealing the change in neuronal activity before ejaculation. Initial thrusts in all MI are accompanied by fluorescence transients, including in the last one (Fig. 5E). The disappearance of the thrust related peaks during the last MI before ejaculation was present in the first session (naive animal) as well as the later sessions (Fig. 5E).

### Optogenetic stimulation of GABA neurons in the VTA interrupts thrusting and mounting behavior

In order to test the functional significance of the VTA DA neuronal activity accompanying mount onset and thrusts, we performed a loss of function optogenetic experiment that consisted in exciting GABAergic interneurons in the VTA, to silence the DA neurons (Fig. 6A). For that we made use of the VGAT-ChR2 mouse line, where the light gated-opsin ChannelRhodopsin (Boyden et al. 2005) is expressed in all GABAergic neurons (Zhao et al. 2011), and implanted an optic fiber above the VTA (Fig6A top)(Tan et al. 2012). The fiber position was analyzed post-mortem (see Methods).

**Figure 6.**
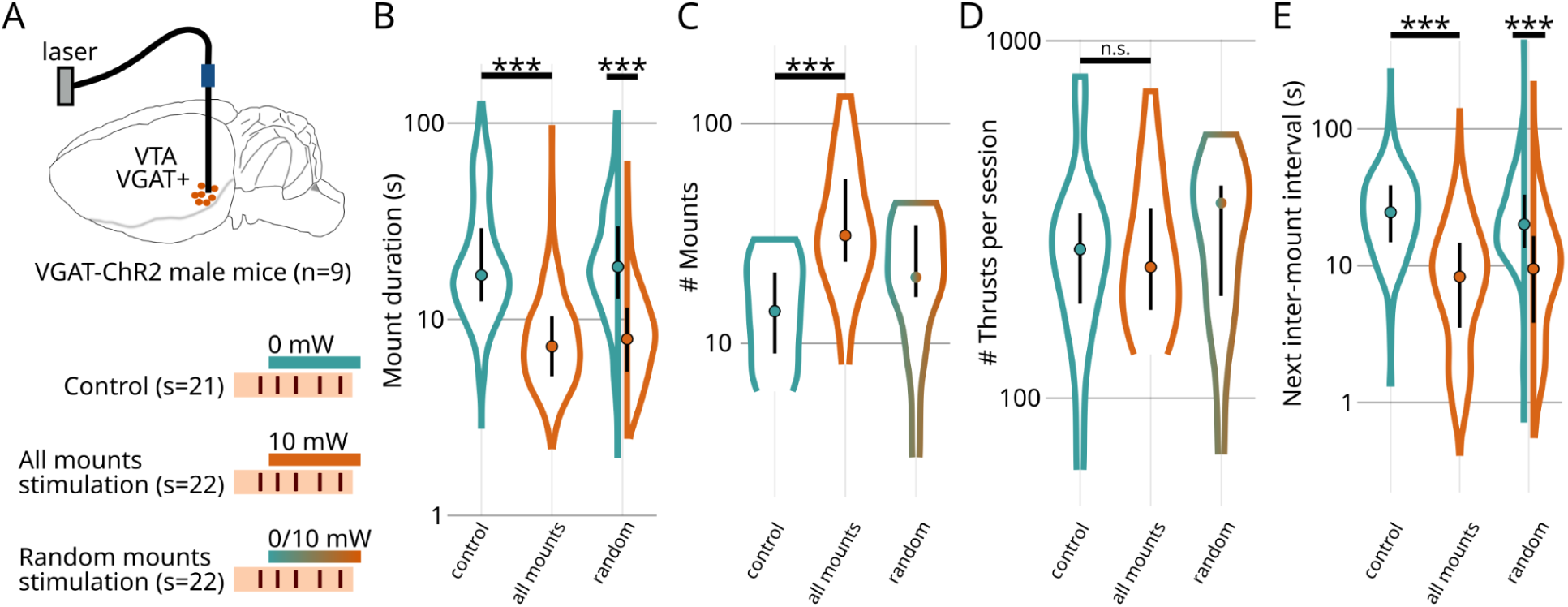
Optogenetic stimulation of GABA neurons in the VTA interrupts thrusting behavior. (A) Optical fiber implantation in the VTA of VGAT-Cre males with ChR2-eYFP (n=9). Three groups of sessions: non-stimulated (Control n=21) and stimulated (All n=22 and Random n=22), after the first Thrust of each MI until they dismounted, with a laser power of 10mW. (B-E) Comparison of the sexual behavior for the non-stimulated (left boxplot, cyan) and stimulated groups (all mounts, middle boxplot, orange; random mounts, right boxplot, cyan/orange for not-stimulated and stimulated mounts): (B) mount (with intromission) duration (in log-scale); (C) number of mounts in the session (in log-scale); (D) total number of thrusts in the session (in log-scale); (E) next inter-mount intervals (in log-scale). Two-tailed Student’s t-test on log-transformed data between control and all mounts and between mounts with and without stimulation for random mounts, ***p<0.001.

We performed light delivery contingent on two different types of behaviors: one when the male was gearing towards the female, which in most cases was followed by a Mount attempt; a second stimulation protocol consisted of turning on the light after the male initiated a MI (after one or two thrusts, by online supervision during the session; the light stayed on until the male dismounted the female). We performed three types of stimulation protocols for each male in the second case: either no MI was stimulated (Control), all MI were illuminated (All) or we delivered light under a random schedule (Random) (Fig. 6A bottom).

Mounting was completely inhibited when we illuminated GABAergic neurons as the animals were approaching the female (Supp Video 1) and only resumed when the light was turned off.

Regarding the illumination once a MI was initiated, we observed that the male stopped thrusting and eventually dismounted the female prematurely. This was the case for the All stimulation group and for the Stimulated MI in the Random group (Fig. 6B). However, interestingly, all animals managed to ejaculate, manipulated or not. We observed that the number of MI needed to reach ejaculation was significantly increased in the All Mounts groups (Fig. 6C). Despite an increase in the number of mounts needed to reach Ejaculation and a reduced duration in the MI duration, on average the three groups of animals performed a similar number of thrusts before reaching ejaculation (Fig. 6D). Interestingly the inter-mount interval was shorter for the All stimulated group and after the stimulated mounts in the Random group (Fig. 6E).

In summary, optogenetically inhibiting VTA DA neurons either prevents mounting behavior before the male attempts copulation, or prevents thrusting if the animals started an intromission, leading to premature dismounting.

## Discussion

In this study we used fiber photometry and optogenetics to investigate the functional importance of VTA DA neurons in male mice. Our study takes a distinct approach by examining how these activity patterns evolve with sexual experience Despite individual variability, the overall structure of mating, multiple sequences of mount-intromission-thrust-dismount, remained consistent across sessions, revealing a stable reproductive pattern. VTA DA neurons were transiently activated when the animals attempted mounting, during thrusts, and ejaculation, and this activity pattern did not change with experience. Notably, as males approached ejaculation, their thrust rate increased while VTA DA activity decreased. Experimental inhibition of these neurons inhibited mounting and disrupted thrusting, provoking early dismounting, but did not prevent ejaculation, suggesting that VTA dopamine signaling provides essential feedback to sustain copulatory behavior.

The VTA-DA activity we observed during mounting, initial penile insertion, and ejaculation is consistent with previously reported findings (Dai et al. 2022; Miyasaka et al. 2025). We found that male behavior remained stable across repeated copulations, and accordingly, the recorded VTA DA transients showed no significant modulation with experience.

Previous studies have examined the effects of sexual experience on mouse copulatory behavior (Jean et al. 2021; Picot et al. 2014). and, similar to our findings, reported changes during the precopulatory phase. However, those studies also observed a reduction in the number of mounts required to reach ejaculation (Picot et al. 2014; Jean et al. 2017), a result not replicated in our work. These discrepancies may originate from methodological differences.

Interestingly, we observed a decrease in VTA dopamine responses to anogenital investigations over consecutive interactions, despite the absence of experience-related changes in VTA DA activity during other phases of sexual behavior. A similar phenomenon has been described in female mice (Moncho-Bogani et al., 2002), where repeated exposure to male pheromones led females to rely less on direct olfactory sampling and more on previously learned associations with contextual cues. This suggests that even in the early, seemingly hardwired stages of sexual interaction, learning mechanisms may be at play, allowing animals to reduce sensory sampling by integrating prior experience. In our case, the diminishing dopaminergic response may reflect a shift from novelty-driven investigation to a learned, expectation-based evaluation of social cues, potentially allowing males to streamline their behavior in familiar contexts. Importantly, as observed in the learning task to which the animals were subjected, the VTA dopamine neurons we recorded are capable of dynamically modulating their activity during anogenital investigations. This demonstrates that these neurons retain functional plasticity in this behavioral context and therefore strengthens our finding that their activity remains unchanged across other components of copulatory behavior.

Sexual behavior is an innate behavior that is usually considered to have intrinsic rewards: ejaculation and possibly thrusts. We decided to use a motivational theory with intrinsic rewards to explain the behavior, namely the homeostatic reinforcement learning (HomeRL) framework developed in (Keramati and Gutkin 2014)(Fig. 7). In this framework, internal rewards are delivered to the subject when an action reduces its homeostatic drives. For sexual behavior, we consider that the animal is subject to two drives: mounting drive and ejaculation potential. The homeostatic point corresponds to no mounting drive and no ejaculation potential. Before the male meets the female, its internal state is at the homeostatic point. The interaction with the female and the arousing cues, like pheromones acquired through anogenital sniffing, make the mounting drive gradually increase. After the mounting drive reaches the mounting threshold, the male attempts to mount the female. If successful (if there is penetration), the male will perform intra-vaginal thrusts. Each thrust has two concomitant effects on his internal state: 1. it reduces the mounting drive (Hull 1966) and 2. increases the ejaculation potential (Fig. 7B). Therefore the male is able to reduce its thrusting drive to zero until dismounting, at the cost of increasing the ejaculation potential. After several mounting–thrusts–dismount sequences, the ejaculation potential is driven close to its threshold. At this stage, a new policy takes over: thrusts do not decrease the mounting drive anymore, only ejaculation can do it, such that the male does not dismount before ejaculating; this way the ejaculation potential keeps increasing until crossing the threshold. We propose that this point of change in policy occurs when emission is triggered, which has also been designated as the commitment state, the point where ejaculation cannot be stopped (Lucio et al. 2011). At this point, removing the penis from the female’s reproductive organ would lead to expulsion of semen in the exterior, an unwanted outcome. Therefore, the change in policy ensures that the male stays mounted and thrusting until the full ejaculatory reflex is triggered. Expulsion occurs only after the male stops thrusting (Lenschow et al. 2025), which is accompanied by the large VTA DA transient. Reaching ejaculation makes the internal state of the animal return to the homeostatic point, associated with a reward.

**Figure 7.**
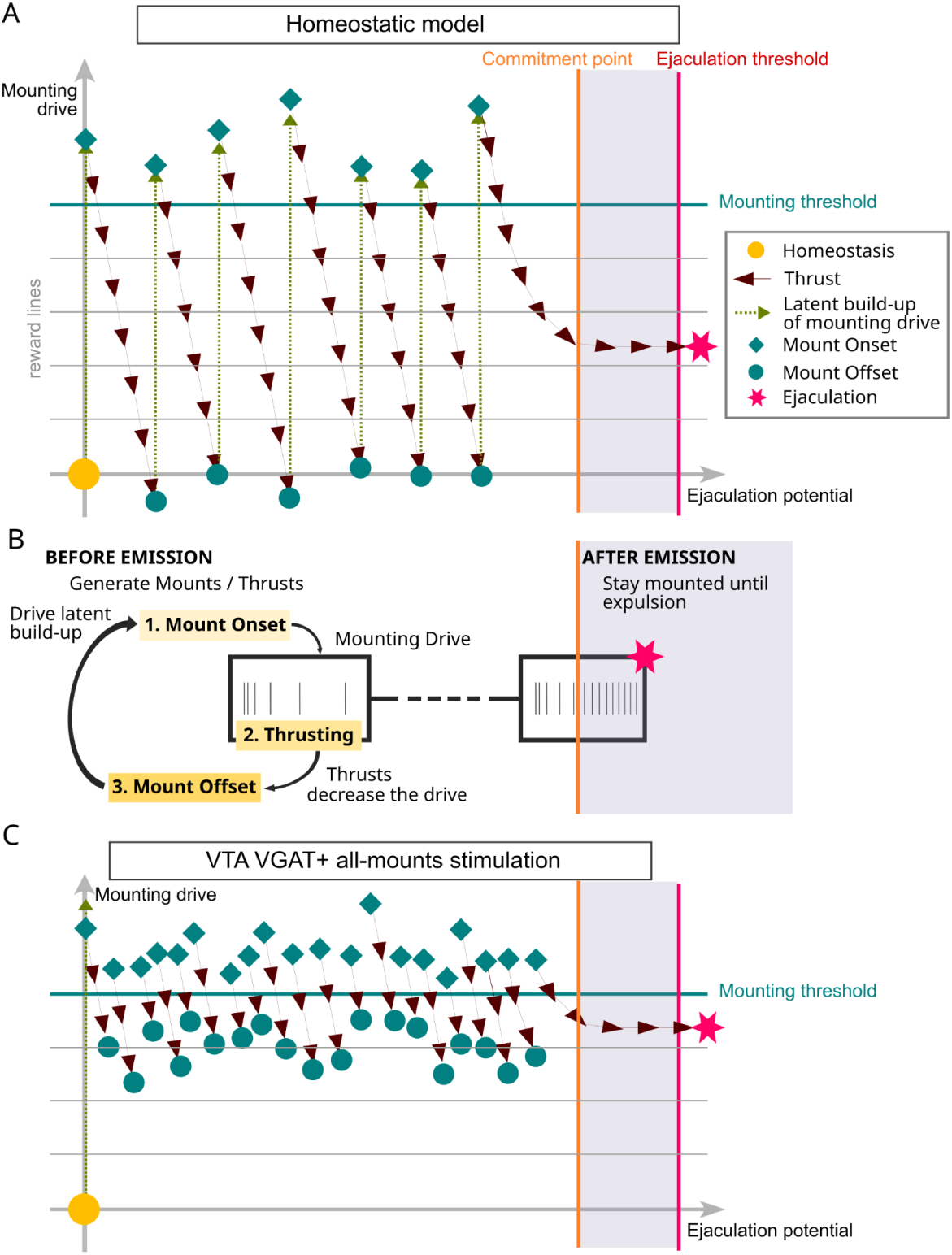
Model of male mice sexual behavior. (A) Homeostatic model of the male sexual behavior. (B) Schematic of the two policies of the male mice during sexual behavior, before and after the commitment or emission point. (C) Homeostatic model of the male sexual behavior with VTA VGAT+ stimulation during all the mounts of each session.

We found that neuronal activity in the VTA was neither associated with, nor predictive of, the progression of sexual behavior. Likewise, no significant changes in behavior were observed until the final mount. Based on these findings, we hypothesize that the neural circuits governing ejaculation and those driving sexual behavior are only loosely coupled. In the proposed model, we distinguish between two largely independent drives: a *mounting drive* and an *ejaculatory potential*. Since the behavioral sequences of mounting and thrusting occur without the need for learning (in fact, multiples rounds of the mounting-thrusting-dismounting are a signature of mouse sexual behavior observed in laboratory and wild derived mice (Valente et al. 2021; Lenschow et al. 2025) and even wild mice (Dewsbury 1975), the dopaminergic activity observed during these events likely reflects the motivation to sustain ongoing behavior. In contrast, the sharp increase in activity immediately following ejaculation, may represent a reward signal reinforcing the overall copulatory sequence.

The optogenetic results support the main hypothesis of our HoRL model, that thrusts have two consequences: a reduction of the drive to mount and an increase of the ejaculation potential (Fig. 7C). In the mounts with stimulation, the male dismounted after a few thrusts, therefore each mount contributed much less to increasing the ejaculation potential than mounts without stimulation, therefore more mounts were needed to ejaculate. Still, the total number of thrusts to reach ejaculation was similar in the two groups, so disturbing the mating pattern did not affect the mechanism to increase the ejaculation potential. At the same time, when the male dismounted, its thrusting drive was still high, so his motivation to mount again the female was high, therefore leading to a reduction of the inter-mount interval. Note that a similar mechanism could be affecting the female and her enhanced motivation to mount again could also contribute to decreasing the inter-mount intervals.

Others have used the same optogenetic strategy (Tan et al. 2012) to disrupt reward consumption and induce place aversion, without notable side effects. In our study, activation of Vgat+ neurons, and the consequent inhibition of VTA dopamine neurons, transiently suppressed ongoing sexual behavior but did not appear to negatively impact male-female interaction. Males readily re-approached the female and resumed mounting immediately after the light stimulation was terminated, and the intermount intervals were even shorter compared to controls. We propose that the shorter intermount interval is caused by the fact that in the stimulated mounts the drive mount was not reduced to zero, so the time to reach the mounting threshold was reduced. This result also points to the lack of an aversive effect with the inhibition of DA neurons.

Intravaginal thrusting relies on the pelvic thrusting reflex—a spinal reflex that is normally under inhibitory control from the brain (Ågmo 2011; Hull and Dominguez 2007). Upon mounting, the male begins thrusting immediately in an attempt to locate the vaginal opening. If penetration occurs, the initial thrusts may proceed independently of brain input, driven primarily by spinal circuitry. This could explain why some thrusting persists even after optogenetic inhibition of DA neurons. However, sustaining thrusting behavior for 10–20 seconds likely requires engagement of higher-order circuits, such as the VTA, as sensory feedback. When VTA DA activity is suppressed, the absence of this feedback may reactivate inhibitory pathways that normally suppress the spinal reflex, thereby preventing continued thrusting.

Regarding mounting-related activity, we observed a ramping signal that began even before the male placed his forepaws on the female or initiated penile insertion. Similar transients have been reported in previous studies during approaches between conspecifics (Gunaydin et al. 2014) and are thought to reflect a motivational signal necessary for initiating social interactions.

In this study, we focused primarily on understanding sexual behavior from the perspective of the male alone. The female’s contribution was not directly examined and was instead treated as a source of variability. However, it is well established that females often control the pacing of copulation to maximize the likelihood of successful fertilization (Johansen et al. 2008). Thus, male behavior may in fact be shaped by the need to follow the female’s pace. Future studies should address this question by including naturally cycling females, as the ovariectomized, hormone-supplemented females used here may display altered behavioral dynamics (Gutierrez-Castellanos et al. 2025).

Together, our results uncover a new functional role for VTA dopamine signaling in male sexual behavior: not as a driver of learning or behavioral progression, but as a feedback signal that sustains ongoing copulatory action. We identify a dissociation between the mechanisms governing the execution of mounting–thrusting sequences and those controlling ejaculation. This separation motivates a conceptual framework in which sexual behavior is organized by interacting but partially independent control processes: a mounting drive that sustains engagement with the female, and an ejaculation potential that accumulates across copulatory bouts. Within this framework, VTA dopamine activity does not encode progress toward ejaculation but rather stabilizes behavior within the appropriate regime, allowing repetition of mounting and thrusting until a separate threshold is reached. Our findings thus argue against a unitary reward-based account of sexual behavior and instead support a control-oriented view, in which motivation, spinal reflexes, and hypothalamic and midbrain circuits cooperate to regulate a structured, evolutionarily conserved behavioral sequence.

---

**Supplementary Table 1.** Sexual behavior performance of individual mice across session 1-6. Schematics showing the performance across time for the ten male mice used in this study. Green tick marks represent sessions where male reached ejaculation, whereas the red cross marks denote sessions where the session was aborted because the male did not try to mount the female. The interval between two consecutive successful sessions (with ejaculation) is one week.

|  | S01 | S02 | S03 | S04 | S05 | S06 | S07 | S08 | S09 | S10 | S11 | S12 | S13 |
| --- | --- | --- | --- | --- | --- | --- | --- | --- | --- | --- | --- | --- | --- |
| #827 | ✓ | ✓ | ✓ | ✓ | ✓ | ✓ | — | — | — | — | — | — | — |
| #892 | ✓ | ✗ | ✗ | ✓ | ✗ | ✓ | ✓ | ✗ | ✓ | ✓ | — | — | — |
| #895 | ✓ | ✗ | ✗ | ✓ | ✗ | ✓ | ✓ | ✓ | ✓ | — | — | — | — |
| #897 | ✗ | ✓ | ✗ | ✓ | ✗ | ✓ | ✗ | ✓ | ✓ | ✓ | — | — | — |
| #986 | ✓ | ✗ | ✓ | ✓ | ✓ | ✓ | ✗ | ✓ | — | — | — | — | — |
| #987 | ✓ | ✗ | ✓ | ✓ | ✓ | ✓ | ✓ | — | — | — | — | — | — |
| #1573 | ✓ | ✗ | ✗ | ✓ | ✗ | ✗ | ✓ | ✗ | ✓ | ✓ | ✗ | ✗ | ✓ |
| #1585 | ✓ | ✗ | ✓ | ✓ | ✓ | ✗ | ✓ | ✗ | ✓ | — | — | — | — |
| #1693 | ✓ | ✓ | ✓ | ✓ | ✓ | ✓ | — | — | — | — | — | — | — |
| #1695 | ✓ | ✓ | ✓ | ✓ | ✓ | ✓ | — | — | — | — | — | — | — |

**Supp Figure 1.**
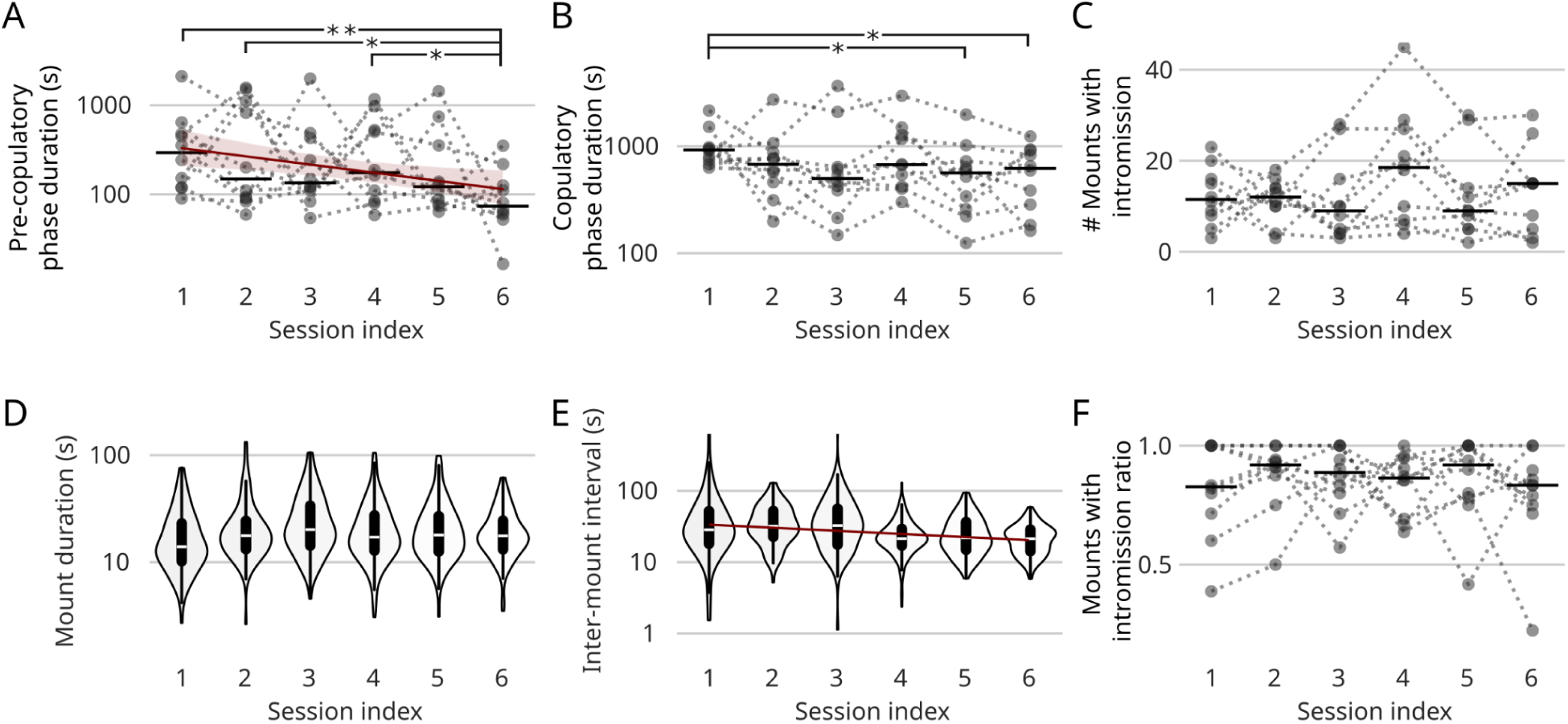
Sexual behavior across sexual experience. (A) Pre-copulatory phase duration (in log-scale) across sessions. Regression line with significant slope, p = 0.01 (R² = 0.11). (B) Copulatory phase duration (in log-scale) across sessions. Regression with non-significant slope, p = 0.06. (C) Number of mounts with intromission across sessions. Regression with non-significant slope, p = 0.5. (D) Mount with intromission durations (in log-scale) across sessions. Regression with non-significant slope, p = 0.2. (E) Inter-mount intervals (in log-scale) across sessions. Regression line with significant slope, p < 0.0001. (F) Ratio of the number of mounts with intromission and total number of mounts across sessions. Regression with non-significant slope, p = 0.7. (A-C) Between sessions, two-tailed Student’s t-test: ***p<0.001, **p<0.01, *p<0.05.

**Supplementary Figure 2.**
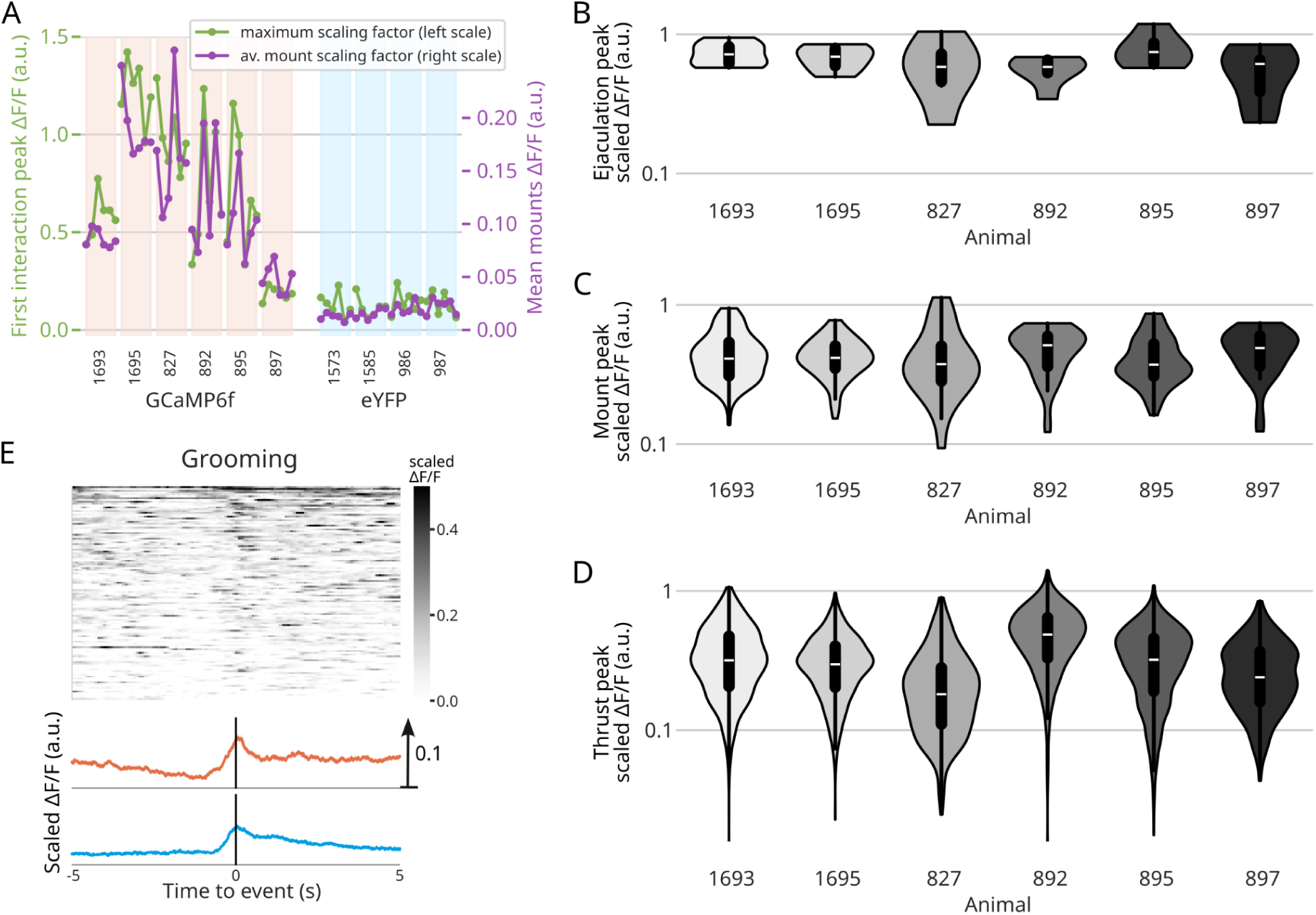
GCaMP signal scaling, GCaMP peak per animal and activity around grooming. (A) Scaling factors for the Ca+ activity signal per session, for GCaMP animals (pink background) and YFP animals (cyan background), comparing the two methods: maximum signal around the first interaction (green dots and lines, scale on the left axis) and average signal during mounts (purple dots and lines, scale on the right axis). (B-D) Peak of activity of the scaled ΔF/F in log-scale across animals for different events: (B) Ejaculation, (C) Mount onset and (D) Thrust (high-pass filtered). After scaling the signal using the peak at the first interaction, the signal amplitude is comparable between animals. (E) (Top) Raster of the scaled ΔF/F for the GCaMP sessions, aligned to Grooming events. The scaled ΔF/F amplitude from 0 to 0.5. (Bottom) Median scaled ΔF/F aligned to the event for YFP sessions (bottom, blue); for the GCaMP sessions (top, red).

**Supplementary Figure 3.**
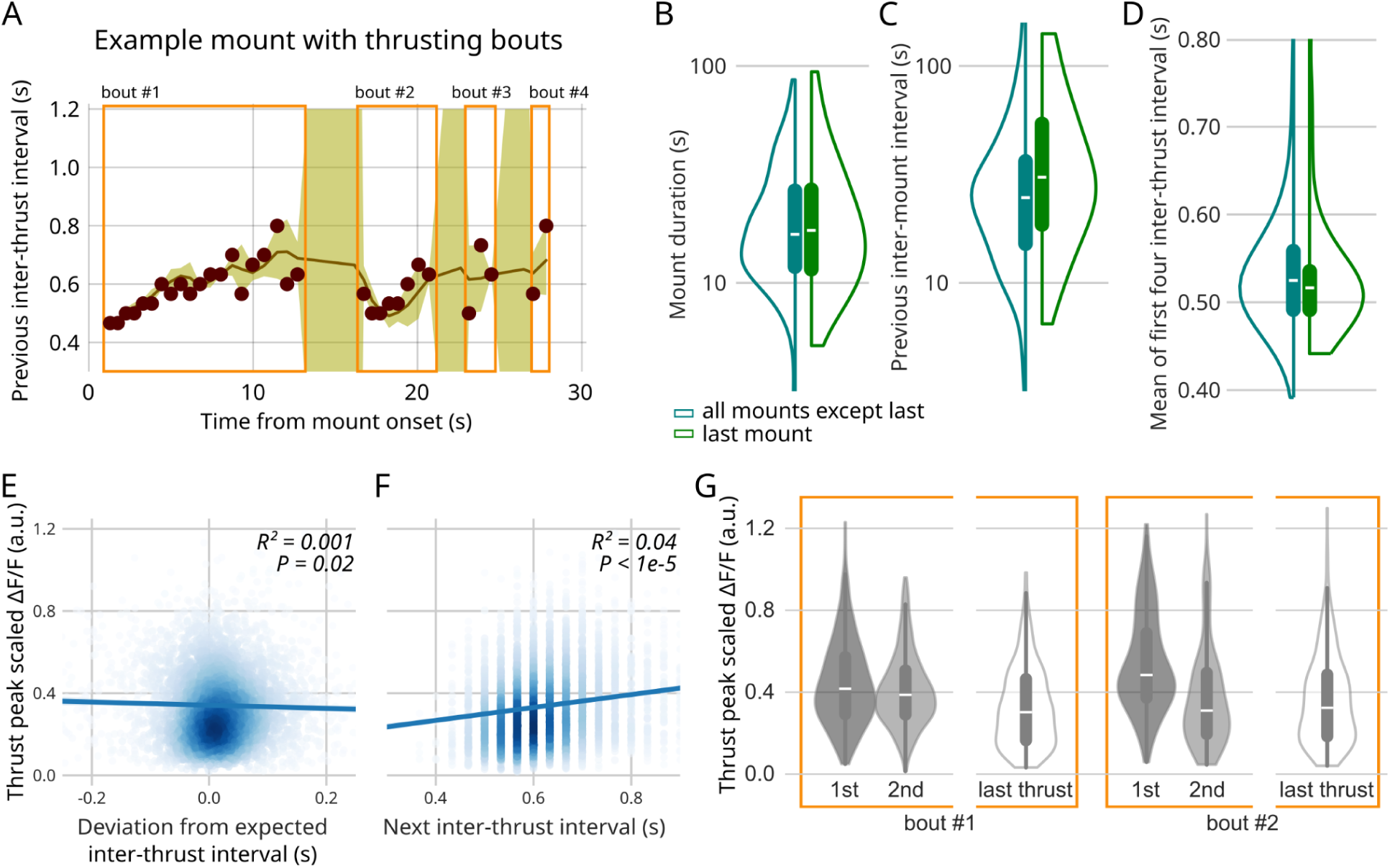
Thrusting bouts, comparison of the last mount with other mounts and thrust peak differences. (A) Thrusting bouts: example mount with four thrusting bouts. Dots show the inter-thrust interval versus time from mount onset. The inter-thrust interval increases inside each bout. The different bouts are delimited by orange rectangles. A new bout starts after an ITI is longer than 1.2s. The expected ITI (brown solid line) is computed by local ridge regressions using the previous 3 to 5 ITIs. Error bar (yellow background): deviation from the expected ITIs. (B-D) Statistics of the last mount (right green violinplot) and other mounts (left cyan violinplot). (B) Mount duration (log-scale), (C) previous inter-mount interval (log-scale) and (D) mean of the first four inter-thrust interval (log-scale). Two-tailed Student’s t-test, all non-significant. (E) Thrust peak (scaled ΔF/F, high-pass filtered) versus next inter-thrust interval. Pearson’s R² and test for the null hypothesis that the correlation is zero. (F) Thrust peak (scaled ΔF/F, high-pass filtered) versus deviation from the expected inter-thrust interval. Pearson’s R² and test for the null hypothesis that the correlation is zero. (G) Thrust peak (scaled ΔF/F, high-pass filtered) for the 1^st^, 2^nd^ and last thrusts of the first bout and for the 1^st^, 2^nd^ and last thrusts of the second bout. The two bouts are delimited by orange rectangles.

**Supplementary Figure 4.**
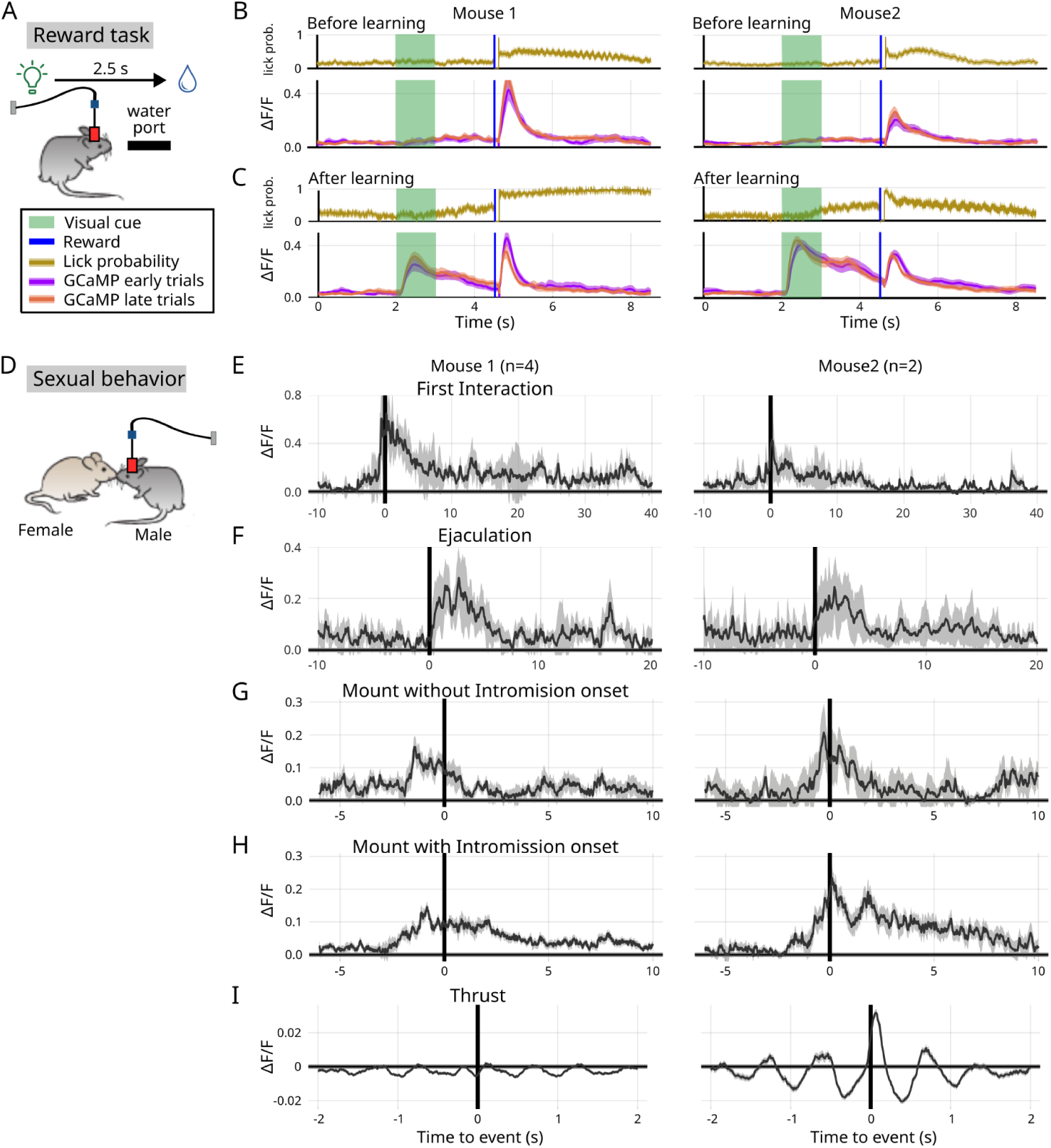
GCaMP signal scaling, other events and reinforcement learning task. (A) Schematics of the classical conditioning setup (B) Probability of licking (top) and photometry recordings of VTA-DAT+ neurons (bottom) during a cue-reward task for 2 animals (2 columns), before learning. The Ca+ activity shows a peak at the reward (blue vertical line) associated with an increase of the licking probability. (C) Same as (B) but after learning the task. The peak at the reward is still present and there is another peak at the visual cue (green background). We observe anticipatory licking after the visual cue. (D) Schematics of the sexual behavior sessions. (E-I) Sexual behavior for the same 2 animals (2 columns) as in B-C. Median (not-scaled) ΔF/F aligned to the events: (E) First interaction, (F) Ejaculation, (G) Mount without thrust onset, (H) Mount with thrust onset and (I) Thrust (high-pass filtered).

## Materials and Methods

### 1. Animals

All experiments were approved by the Champalimaud Centre for the Unknown Bioethics Committee and by the Portuguese Veterinary General Board and in accordance with the European Union Directive 86/609/EEC.

All animals were bred in-house at the vivarium of the Champalimaud Research department. All male mice had the DAT::Cre genotype (Jax 006660), and females were C57BL/6J (Jax 000664). For the optogenetics experiments we use Vgat-ChR2-YFP males (Jax 014548). All animals were maintained on a 12h light/12h dark cycle.

Adult (12 weeks of age onwards) male and female mice were used. Male mice were single housed after 8 to 12 weeks of age, whereas female mice which were subjected to ovariectomy surgery between 8-12 weeks of age, were group housed and underwent regular oestrogen and progesterone injections in order to induce sexual receptivity (Gutierrez et al 2025). Female mice were also sexually trained with male studs prior to being used in the experiments.

### 2. Viral injection and optical fiber implant

Surgical procedures were performed under oxygen and isoflurane mixture anaesthesia (1-2% at 1L/min) and controlled animal body temperature. Each animal was given two unilateral viral injections at VTA using a stereotaxic system (Kopf) with the following coordinates: -2.87 anteroposterior, +/- 0.65 lateral from bregma and -4.3 and -4.7 deep from the brain surface (REF). The injections were done through a micropipette pulled from borosilicate capillaries (Drummond Scientific Company) and using an automated microprocessor (Nanoject II, Drummond Scientific Company) at a 0.1 Hz frequency pulses, with 4.6 nl injection volume per pulse, at a total volume of 330 nl of AAV5.Syn.Flex.GCaMP6f.WPRE.SV40 (U.Penn, Vector Core) or 125 nl of AAV5.EF1a.DIO.eYFP.WPREpA (UNC, Vector Core) per site. For all injections, the micropipettes were left for 5 min at the VTA coordinates before starting injecting the virus and for 5 min after the last pulse, before withdrawal of the needle. We chronically implanted optical fibers with 6 mm long and 400 μm of diameter (MFC 400/430-0.48 6mm SM3 FLT, Doric Lenses) in VTA coordinates: -2.87 anteroposterior, +/- 0.65 lateral from bregma and -4.5 deep from the brain surface. Immediate post operative care included iodine and wound powder around the implant, and food supplemented with analgesic (MediGel, 1 mg carprofen/2 oz cup) for the first 2 days post-op.

The same procedure was performed in order to implant an optic fiber above the VTA for the VGAT-ChR2-YFP males.

### 3. Sexual behavior

All behavior experiments were performed during the night (mice were kept in reverse light cycle conditions, 8am-8pm).

The behavioral arena was custom made and consisted of a 35.5 × 13.4 × 21.5 cm box with transparent acrylic walls, two fixed point grey cameras (FL3-U3-13S2C-CS) and uniform and indirect illumination which was provided by one set of infrared led lights placed around the arena.

The male was first placed in the behavioral arena and after a random period of habituation time to the box, a sexually experienced and receptive female mouse was introduced in the opposite place where the male was currently standing. The two animals were left to behave freely, until the male ejaculated and the female was removed from the box. If after 1h, the male did not attempt to mount the female, the session was prematurely terminated. After a session with ejaculation, the time limit for waiting for a mount attempt was reduced to 30 min. After each session, regardless of the ejaculation outcome, the male was given one week before being tested again.

We allowed 10 sexually naive male mice to sexually interact with receptive females during 6 sessions where the male ejaculated. Other sessions, when the male or the female did not engage in sexual behavior were discarded. The stimulus females were different for each session, except for the sessions 4 to 6, where the female was the same to assess the variability due to the female in the sexual behavior.

### 4. Sexual behavior annotations

All the sessions of sexual behavior were recorded with two 30 fps cameras recording from above and from the wider side of the box. The videos were later manually annotated on a custom made Bonsai workflow (Lopes et al., 2015) to identify the time of the following sexual behaviors. For the appetitive phase: time of First interaction between the male and the female, and male performing Anogenital investigation to the female. For the consummatory phase: Failed mount attempt, Male mounting the female (the Mount onset and Mount offset times were annotated), intra-vaginal Thrusts and Ejaculation. Mount onset corresponds to the male putting his paws on the flank of the female, with his body aligned with her. The Mount onset was exactly annotated when the two bodies became aligned. Mounts can have penile insertion and Thrusts or not. Thrusts were precisely annotated when the male pulled out the penis and his pelvis was furthest away from the female body.

We also analyzed the inter-Mount intervals (IMI) that are defined as the time between the previous Mount offsett and the next Mount onset. Likewise we defined the inter-Thrust interval (ITI) as the time between two consecutive Thrusts events. The mating time, from first interaction with the female to ejaculation, was divided into the latency to first mount, where the first mount could be with or without penile insertion, and the ejaculation time, the time from first mount to ejaculation.

For analyzing the thrusting rhythm, we only take into account the thrusts from the first thrusting bout. Thrusting bouts are defined by sequences of thrusts with inter-thrust intervals shorter than a threshold that we define below. When an ITI is longer than this threshold, the next thrust belongs to a new thrusting bout. The threshold was set to 1.2s as it corresponds to twice the median ITI, a robust estimate of the average ITI.

### 5. Classical Conditioning Setup

The behavioural box (10 × 10 × 10 cm, custom made) contained one central nose port with an infrared beam emitter/sensor to measure nose poke in and out, a solenoid valve (LHDA1233115H; The Lee Company, Westbrook, CT) to deliver the reward and a yellow LED light to act as the conditioned stimulus (CS). We added a custom made sensor to measure the reward’s consumption in the form of a lick-o-meter. Sensors were accessed using a microcontroller (Arduino Duemilanove). All electrical signals were recorded with a National Instruments data acquisition card. The task was implemented by a desktop running a custom Bonsai (Lopes *et al*., 2015) state machine software through the microcontroller.

**Figure 1.**
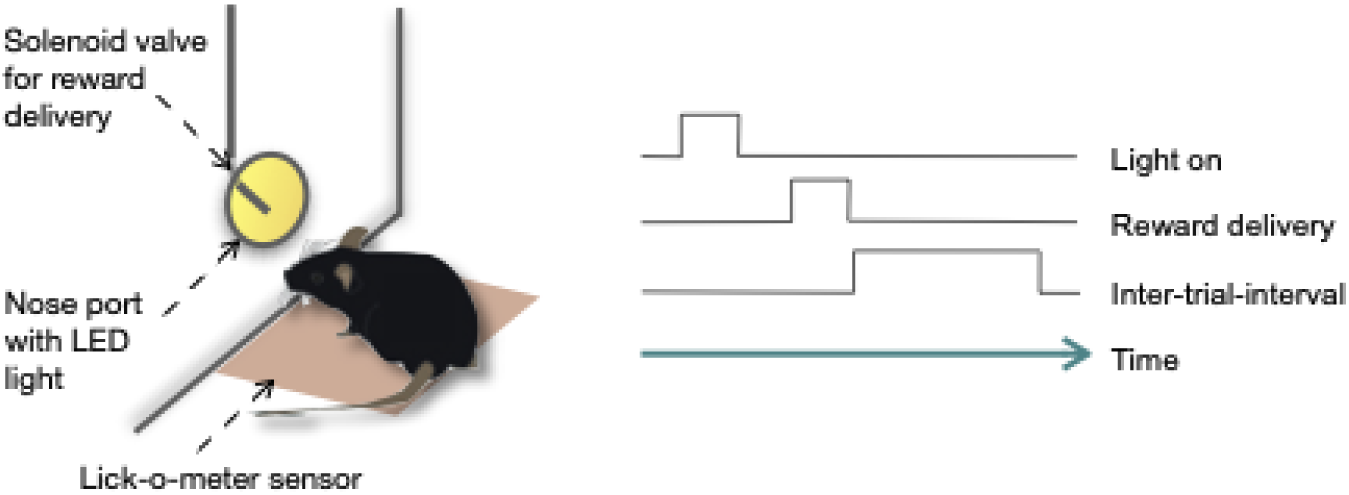
Scheme of the classical conditioning setup and trial structure. Schematics of the classical conditioning setup (left) and order of events within a trial (right).

### 6. Classical Conditioning Task

To avoid a decrease in the motivation to engage in the task, male mice were water and food-deprived for 24h before performing the task. Each session had a maximum length of 60 minutes. Trial onset was prompted by the onset of the yellow LED light for 1 second. After the light was on, mice were able to collect a reward (approximately 6µL of condensed milk diluted 1/10 in water) before a new trial started (after a randomised inter-trial-interval that ranged from 5 to 42 seconds). The task setup and structure is depicted in the figure below.

### 7. Fiber photometry

LED fiber light source (465 nm; LED-FLS 465, Doric Lenses) was coupled to individual patch cords (100 μm core diameter, 0.22 NA, Doric lenses) and connected to an individual collimator adapter each (EFL 4.5 mm, NA 0.50, Doric lenses) and a neutral density filter (NE30B, Thorlabs), mounted on the main setup unit. Two dichroic mirrors (Di01-R561-25x36, Semrock, T495LP, Chroma) were fixed inside the main unit, allowing for the 473 nm light delivery and collection of light emitted by GCaMP6f or eYFP. The 473 nm light was coupled into a patch cord (200 μm core diameter, 0.48 NA, Doric lenses) using a lens (EFL 4.5 mm, NA 0.50) and a rotatory joint (Doric lenses). The patch cord was connected to the animals’ chronically implanted optical fiber (400 μm core diameter, 0.48 NA; MFP 400/460/900-0.48 1m FC-CM3, Doric lenses). For detection of GCaMP6f or YFP fluorescence, light was collected by the lens, transmitted and reflected by the dichroic mirror before final filtering and focusing (bandpass: ET525/50m; Chroma, lens: EFL 25.4 mm, model LA1951-A, Thorlabs) into a photodetector (PDF10A/M, Thorlabs). Photodetector output was digitized at 4 kHz (NI 6351 board) and recorded with custom Bonsai (Lopes et al., 2015) software. A custom made base station and HARP Software (Champalimaud Foundation Hardware Platform) were used in order to control and trigger the synchronized acquisition of the videos and the fluorescent signals from the photodetector.

### 8. Optogenetic experiment

Males implanted with an optic fiber over the VTA were allowed to have several rounds of sexual interaction with OVX females injected with sex hormones to induce receptivity.

Blue light to excite ChR2 was turned on manually either when the male was approaching the female or after he achieved penile insertion and executed 1-2 thrusts. When the stimulation was performed during the MI, light was either delivered on All mounts, randomly or never (control session). Illumination was delivered at 10mW.

### 9. Processing of the fluorescence signal

All the data analysis was performed with custom code written in Julia.

Raw photometry traces were first down-sampled from a sampling rate of 4 kHz to 100 Hz. For each session we calculated the ΔF/F as (F - F_0_) / F_0_, where F_0_ was computed as the 20% quantile of a 3 minute rolling window preceding each time point.

To normalize ΔF/F to peak at the first interaction after Female In, the normalization factor was computed as the maximum signal in a 20 s window around this first interaction event.

To normalize ΔF/F to the average mount signal, the normalization factor was computed by taking the mean ΔF/F for each mount (taking 1s before the onset and 1s after the offset), then taking the mean of all the mounts.

Normalizing by the peak at the first interaction is not possible for eYFP animals as there is no such peak. However, to make them comparable to the GCaMP6f animals, we divided the eYFP signals by the average normalization factor from the GCaMP6f animals, which was equal to 0.5.

For the purpose of computing peri-event time histograms, the ΔF/F signal was filtered to remove the low-frequency components with a forward-backward highpass Butterworth filter of order 8 and zero phase with cutoff frequency of 0.8 Hz, which is half the median inverse inter-thrust-interval.

Rasters of peri-event ΔF/F were ordered by the maximum signal amplitude in the window of interest.

Two different algorithms were used to compute the amplitude of the ΔF/F peaks:

- for the mount onset peaks: the ΔF/F was smoothed using a Gaussian filter with a standard deviation of 50 ms (5 time frames). Then we determined the local maximum of the smoothed ΔF/F in an interval of 2 s before and after the annotated mount onset event; the higher bound of this interval was cropped to 0.5 s before the first thrust, if the first thrust happened before 2.5 s.
- for the thrust peaks: all the peaks at thrust events were modeled by pulses with the same shape, namely a double-exponential or alpha function with a fast rising time of 50 ms and a decay time of 150 ms, which is compatible with the values found for GCaMP6f dynamics (https://doi.org/10.1038/srep38276). To account for the overall increase of ΔF/F after mount onset, instead of fitting to ΔF/F, the highpass filtered signal *z* was used, relative to its lower envelope so the fitted signal was always positive. The lower envelope *b* was defined as the solution of a constrained quadratic optimization problem, *L_1_ = (z – b)² + η (b_i+1_ – b_i_)²*, (η=10^5^) with the constraint that *z >= b*. Then, the relative signal *y = z – b* was fitted to a sum of pulses, having exactly one pulse for each annotated thrust event *t_i_*. The amplitudes *a_i_*of the pulses and the delays *τ_i_* from the annotated events were fitted using a quadratic loss function with quadratic regularizations for both the amplitudes and the delay: *L_2_ = (y – S * a)² + λ |a|² + μ|τ|²* (where *S* was a matrix of single pulses at times *t_i_+τ_i_*). Only the signal during mounts was used to compute the loss function.

### 10. Statistical analysis

One-way analysis of variance (ANOVA) for repeated measurements was used to compare two groups or linear regression if the number of groups was more than 2, using *t*-test for significance test. For positive variables, the data was log-transformed before performing the linear regression or the ANOVA, as they appear in the figures.

Correlation between two variables was estimated by performing a linear regression with intercept of one variable versus the other, computing the r² value and an F-test versus the null model.

Violin plots are truncated to the extrema values, the dot corresponds to the median and the black vertical lines extend up to the first and third quartiles.

In order to compare sessions with different numbers of mounts, we used the so-called session progression, that represents the mount index normalized to the total number of mounts in the session.

Identically, we used mount progression to normalize mounts with different number of thrusts, that corresponds to the thrust index in the mount, normalized to the total number of thrusts in this mount. If only the first thrusting bout is considered, the mount progression corresponds to the thrust index in the mount, normalized to the number of thrusts in the bout.

For plotting various quantities versus mount progression, we used two-dimensional kernel density estimation and the density was fitted with non-parametric conditional kernel density estimation [REF Rosenblatt 1969], that is taking the conditional mean value along the y-axis quantity for each mount progression value.

If not stated otherwise, peri-event time histograms are summarized using the median, where the error bars correspond to the standard error of the median, computed with a sparsity measure of the quantile.

For the correlation of the thrust peak with the expected inter-thrust interval (ITI), we need to estimate an expected ITI. For that we run local ridge regressions using between 3 and 5 consecutive ITIs, the ridge penalty ensures that the slope for these 5 ITIs is not very different from the slope estimated from the previous 5 ITIs. These local models of 5 ITIs are used to predict the next ITI, that we use as the expected ITI. The deviation with the expected ITI is the difference between the observed and the expected ITI. See Supplementary Figure 3 for an example mount with the expected ITIs and the deviation.

### 11. Immunohistochemistry and microscopy

Histological analysis was performed after all experiments to confirm optical fiber placement and viral expression. Mice were anesthetized with pentobarbital (Eutasil, 50 mg/kg intraperitoneally, CEVA Santé Animale, Libourne, France) and perfused transcardially with 4% paraformaldehyde (P6148, Sigma-Aldrich). The animals were then decapitated and the heads left overnight in 4% PFA before removing the brains from the skull and keeping them in 30% sucrose solution with 0.1% azide, for approximately 1 week until sectioning. A sliding microtome (SM 2000 R, Leica Biosystems, Wetzlar, Germany) was used to section the brain into 45 μm thick slices that were then immunostained with antibodies against tyrosine hydroxylase (22941, 1:5000, ImmunoStar) to visualize dopaminergic neurons. Images were acquired with a wide field microscope (AxioImager M2 wide field fluorescence microscope, Zeiss) using a 10X objective.

